# Integrated Field’s metal microelectrodes based microfluidic impedance cytometry for cell-in-droplet quantification

**DOI:** 10.1101/278069

**Authors:** Jatin Panwar, Rahul Roy

**Author notes:** Correspondence: Rahul Roy, Department of Chemical Engineering, Indian Institute of Science, Bangalore, Karnataka, India 560012.

## Abstract

Microfluidic impedance cytometry (MIC) provides a non-optical and label-free method for single cell detection and classification in microfluidics. However, the cleanroom intensive infrastructure required for MIC electrode fabrication limits its wide implementation in microfluidic analysis. To bypass the conventional metal (platinum) electrode fabrication protocol, we fabricated coplanar ‘in-contact’ Field’s metal (icFM) microelectrodes in multilayer elastomer devices with a single photolithography step. Our icFM microelectrodes displayed excellent and comparable performance to the platinum electrodes for detection of single erythrocytes with a lock-in amplifier based MIC setup. We further characterized it for water-in-oil droplets generated in a T-junction microfluidic channel and found high sensitivity and long-term operational stability of these electrodes. Finally, to facilitate droplet based single cell analysis, we demonstrate detection and quantification of single cells entrapped in aqueous droplets.

## 1. Introduction

Microfluidics allows precise control over chemical and biological investigations at micron scales that are comparable to the dimensions of the biological cell. As a result, microfluidic detection and analysis of single cells has seen a tremendous rise (Mehling and Tay 2014; Reece et al. 2016; Streets et al. 2014) and has enabled lab-on-chip and point-of-care applications (David J. Beebe et al. 2002; H.A. Stone et al. 2004; Jung et al. 2015; Shi et al. 2012). In many microfluidic applications, cell detection is achieved using optical methods. Sensitive methods like fluorescence can allow molecule-specific detection enabling cell classification. However, these methods require additional and sometimes tedious labelling steps (Specht et al. 2017). Therefore, there have been parallel efforts to develop label-free assays for the detection and characterization of cells (Blasi et al. 2016; Kim et al. 2017; Shapiro 2005; Watson 1991). One such promising technique for single cell analysis is Microfluidic Impedance Cytometry (MIC) which relies on cellular electrical impedance (Holmes and Morgan 2010; Hywel et al. 2007). The dielectric properties of a biological cell are defined by cellular characteristics like cell volume, composition and architecture. Continuous flow microfluidic devices with embedded microelectrodes for electrical measurements can be employed for detecting and classifying single cells in a high throughput manner (Coulter 1956; Han et al. 2012; Hywel et al. 2007; Sun and Morgan 2010; Watkins et al. 2013). In most MIC devices, as the cell suspended in an electrolyte flows over an array of metal electrode pairs connected to a high frequency AC excitation source, the dielectric properties of the media between the electrodes change. This in turn attenuates the measured impedance at a particular frequency across the microelectrode pair and generates a peak in the differential output. Highly sensitive MIC based detection has already shown promise in characterization of sub-cellular features (Hywel et al. 2007; McGrath et al. 2017; Sun and Morgan 2010).

However, MIC devices rely on cleanroom intensive techniques like metal sputtering/vapour deposition to generate metal electrodes (100-200 nm thick) on the substrates (glass or silicon wafer). The substrate is then bonded to the flow channel to complete the assembly of a coplanar microelectrode integrated microchannel (Chidsey et al. 1986; Kannan et al. 2016; Temiz et al. 2012). This elaborate and costly microelectrode fabrication process has limited the deployment of MIC in diagnostics and other real-world applications.

There have been several prior efforts to develop cheaper and efficient embedded microelectrodes. For example, fusible alloy or liquid metal filled into dedicated microchannels and placed in proximity with flow-channels have been employed as non-contact electrodes (Sciambi and Abate 2014; Thredgold et al. 2013). However, these electrodes have limited impedance sensitivity because of the attenuated charge density and electric field strength between the electrodes with the intervening elastomer. As impedance sensitivity has a strong dependence on electrode dimensions and displays optimum sensitivity for electrode widths comparable to the particle size, (Clausen et al. 2015a; Gawad et al. 2004) fabrication of microelectrodes and their placement in close proximity to flow channels is critical for MIC. To address this, alternate architecture(s) where the electrodes are in direct contact, namely, ‘in-contact’ with the flow channel are employed (Guler et al.; Richards et al. 2012; So and Dickey 2011; Song et al. 2011). However, the sensitivity and stability of such microelectrodes for impedance detection of cells and their performance relative to metal electrodes has not been carefully characterized previously.

In this work, we present a simple fabrication method and characterization of MIC compatible coplanar ‘in contact’ Field’s metal (icFM) microelectrodes. Importantly, our icFM microelectrodes displayed impedance signal strengths and contrast that is comparable to the conventional platinum electrodes for single erythrocyte detection in a custom-built suction driven continuous flow setup.

After establishing the stability of icFM microelectrodes and their compatibility with MIC, we used the setup for single cell quantification in droplets, a requirement for many single cell isolation and analysis platforms (Joensson and Andersson Svahn 2012; Rosenfeld et al. 2014; Zhang et al. 2017). Our method provides a cheap, sensitive, non-optical and label-free approach to detect and quantify cells within water-in-oil microdroplets (Lu et al. 2017).

## 2. Materials and Methods

### 2.1 Electrode fabrication

Microchannels for the flow and the microelectrode layers were designed and fabricated using standard soft photolithography methods (Duffy et al. 1998; Thompson 1983). Briefly, microchannels were designed using CleWin 4 and the chrome mask was etched using a mask writer (Heidelberg µPG 501) and EVG 620 (EV Group) was used for UV exposure. Polydimethylsiloxane or PDMS (184 Sylguard, Dow Corning) was used for all the device fabrications. SU8 2015 (MicroChem) was spin-coated on a silicon wafer at 2100 rpm to obtain a thickness of 20 µm prior to UV exposure and development. A PDMS cast of the electrode layer (L1, **∼** 4 mm) consisted of three independent 100 µm channels that reduced to a width of 30 µm and an inter-electrode gap of 20 µm at the detection region (**Figure 1a**). L1 also incorporates all inlet and corresponding outlet ports for molten alloy flow as well as fluid flow. A flat (unpatterned) and thin PDMS sacrificial base layer (L2, **∼** 1 mm) was placed under L1 to cover the channels while taking care to remove all trapped air bubbles between the elastomer layers. The flexibility and low surface roughness of polymerized PDMS provided an excellent seal between the layers. The L1+L2 assembly was placed on a hot plate at 130 °C for 20 minutes. 15 µl of molten FM (Bombay metal house) at 130 °C was pipetted into the electrode channel inlets while the assembly was placed on the hot plate. Using a 50 ml syringe, a suction pressure (**∼** 80 kPa) was applied to the other end of the channel until the metal reached the outlet port for each electrode channel. Suction pressure drives the molten alloy flow through the microchannels as well as holds the two layers together without any additional requirements for bonding. The assembly was allowed to cool after scraping off the excess metal till the FM solidified and the L2 was peeled off to expose the microelectrodes (**Figure 1a**). The flow layer (L3, **∼** 1-4 mm) consisted of a 20 µm high and 70 µm wide channels converging to a 20 µm width at the detection region and runs orthogonally to the electrode channels. In case of water-in-oil droplet measurements, a T-junction was added to the flow channel upstream of the detection region. L1 and L3 were aligned using the guide marks and bonded after a plasma treatment. The bottom layer (L3) was also plasma bonded to a glass slide for device rigidity and ease of operation. To overcome the reduced hydrophobicity of PDMS after plasma treatment, the device was kept overnight in an oven at 45 °C (but below the FM melting point, **∼** 60 °C). The device thus fabricated, contains a fluidic channel passing orthogonally under an array of three ‘in-contact’ coplanar microelectrodes (**Figure 1b**).

**Figure 1:**
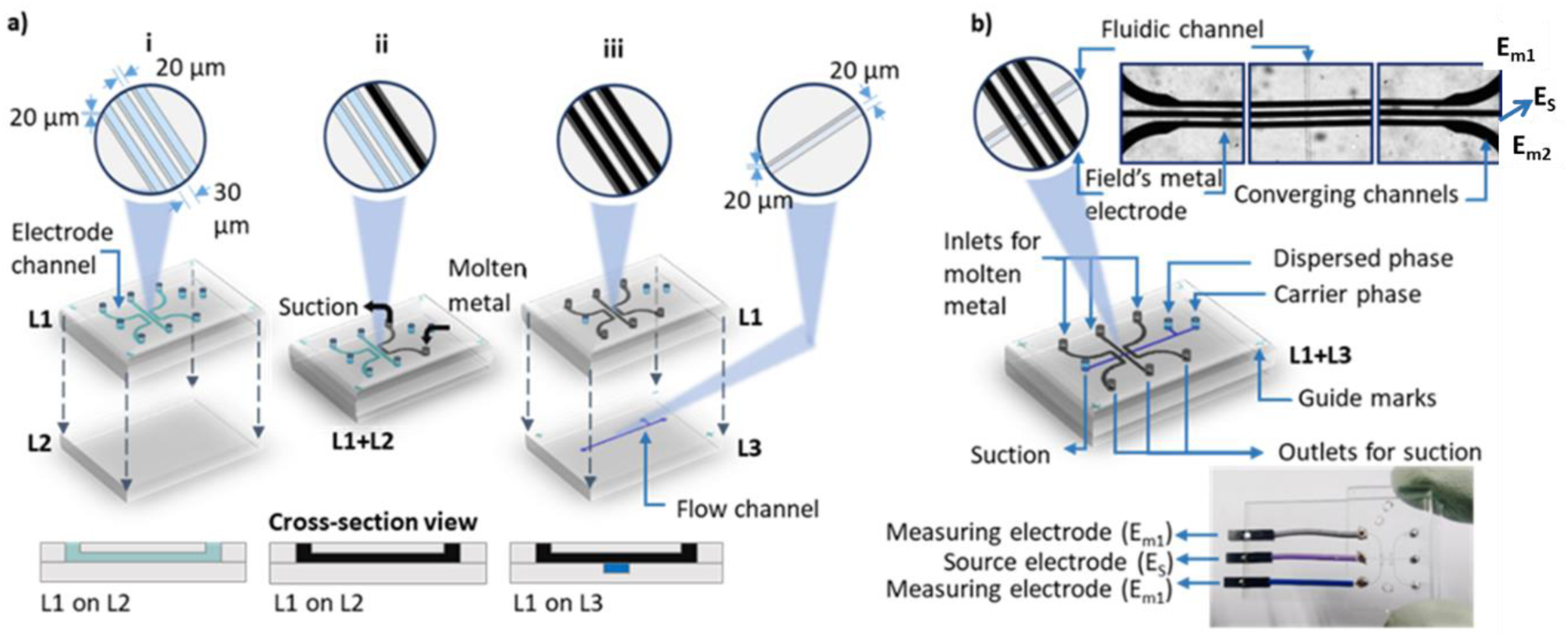
Coplanar ‘in-contact’ Field’s metal (icFM) electrode fabrication workflow **a)** Schematic of fabrication process; **i**: The PDMS layer containing electrode channels (L1) is placed on a flat PDMS sacrificial layer (L2) which acts as a base during electrode casting. **ii**: Molten FM is flown into the electrode channels using suction. L2 is peeled off once the electrodes solidified. **iii:** Electrode layer (L1) is then plasma bonded to the flow layer (L3) **b)** Schematic and optical image (top right) of the microelectrode array and assembled microfluidic device with embedded coplanar icFM microelectrodes is shown.

### 2.2 Fluid flow control

Suction pressure was stabilized using a custom-built active-feedback pressure control module. A syringe pump (1010X, New Era) operation is regulated with a NI USB 6008 data acquisition card and MP3V5050V pressure transducer (**Figure 2a**). A LabVIEW based user interface was designed to control and monitor the applied suction pressure in real-time. The control module facilitates application of suction microfluidics (Abate and Weitz 2011) in long-term operations of multi-phase microfluidic devices and thus reduces the control requirements.

**Figure 2:**
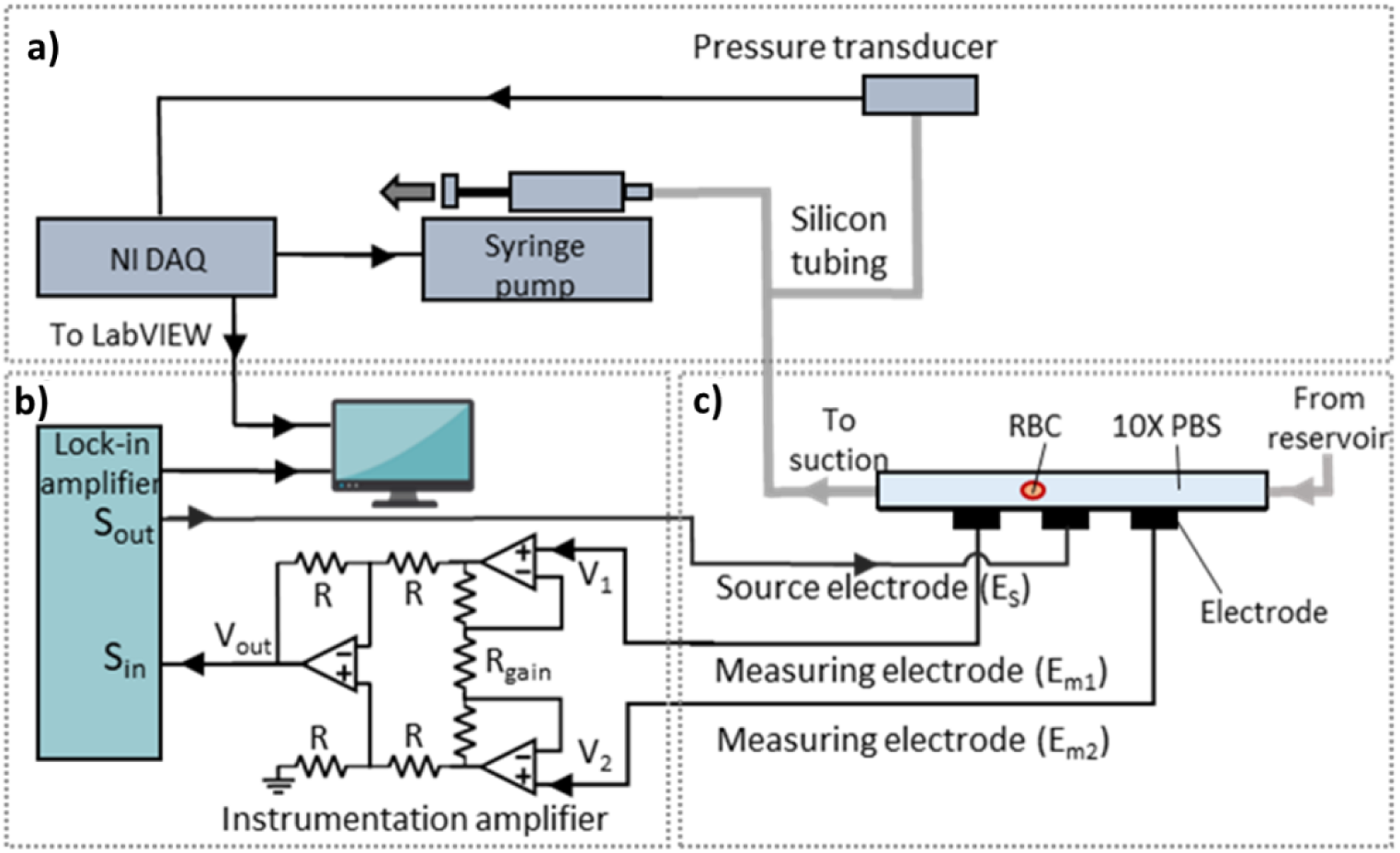
Lock-in amplification based microfluidic impedance cytometry (MIC) setup with feedback controlled suction flow **a)** Schematic of active feedback pressure control module that regulates suction for fluid flow in the microfluidic channel. **b)** Lock-in amplifier with built-in signal generator provides the excitation AC voltage (S_out_) to the source electrode and measures the differential voltage signal (S_in_) from the measuring electrodes via an instrumentation amplifier. **c)** A cross section of the ‘in-contact’ coplanar electrodes orthogonal to the flow channel with its fluidic and electronic connections is shown.

For cell-in-droplet experiments, water-in-oil droplet was generated using a T-junction flow channel architecture (Xu et al. 2008). The continuous phase consisted of fluorinated oil (Bio-Rad, 1863005) with an erythrocyte cell suspension in 1X Phosphate-buffered saline or PBS (Sigma-Aldrich) as the dispersed phase. The capillary number for the continuous phase and Reynolds number at the T-junction were 8.24 × 10^−2^ and 0.15, respectively. Average droplet volume was ∼ 150 picoliter that was generated at a frequency of ∼ 10 Hz using 18 kPa suction pressure at the outlet port.

### 2.3 Microfluidic impedance cytometry setup

A lock-in amplifier with a built-in signal generator (HF2LI, Zurich Instruments) was connected to the microelectrodes via a custom-made low noise instrumentation amplifier circuit. We used the commonly employed MIC electrode architecture: a three coplanar electrodes configuration within the microfluidic chip representing two electrode pairs (Clausen et al. 2015b; Errico et al. 2017; Gawad et al. 2001). An excitation AC signal was applied to the source electrode (E_s_) via the signal-output port of the lock-in amplifier. The two measuring electrodes (E_m1_ and E_m2_) were connected to the instrumentation amplifier circuit (**Figure 2b**).

Particles passing over the electrodes perturb the dielectric properties (and hence the impedance) of the sample between the electrodes (**Figure 2c**). This change is measured as a voltage offset across the two electrode pairs (ΔV = |V_2_ - V_1_|). The instrumentation amplifier converts the output voltage (V_1_ and V_2_) from electrode pairs into a differential voltage (V_out_) with a gain, which was then fed into the signal-input of the lock-in amplifier. ZiControl software was used to record the amplified differential voltage (Vrms) from the lock-in amplifier.

### 2.4 Imaging and analysis

The devices were imaged using an inverted microscope (RAMM, Applied Scientific Instrumentation). Images were acquired using μManager 2.0 (Edelstein et al. 2014). Scanning electron microscope (Ultra 55 FE-SEM, Carl Zeiss) was used for electrode surface micrographs. All the data analysis was done using custom codes in MATLAB.

### 2.5 Sample preparation

The blood samples were provided by the author and drawn by trained personnel at the Health center, IISc Bangalore. For erythrocyte enrichment, 5 ml of blood was spun at 500 x g for 7 mins. The supernatant was discarded and the pellet was resuspended in 5 ml of 10X PBS (or 1X PBS in case of droplets). The samples thus prepared were visually inspected on a microscope to ensure single cell suspension (**Supplementary figure S1**).

## 3. Results and discussion

Our MIC microfluidic module employs a single standard photolithography workflow to generate two PDMS microchannel layers, i.e. the microelectrode layer (L1) and the flow layer (L2) (**Figure 1**). First, the electrodes are fabricated separately by filling the L1 microchannels with a fusible alloy (FM). The channel architecture helps define the arrangement, dimensions and the position of the microelectrodes. We chose FM (32.5% Bi, 51% In, 16.5% Sn) among the eutectic fusible alloys because of its non-toxic nature. FM also has a melting point of **∼** 60 °C which is significantly above room temperature. The flow and electrode layer (with electrodes placed orthogonal to flow channels) are bonded to assemble the complete icFM device in a cleanroom-free environment thus significantly reducing processing time and resource requirements (**Supplementary table T1**). Deployment of thus fabricated icFM microfluidic device in a MIC setup with an active feedback pressure control module is described in the Methods and **Figure 2**.

### 3.1 icFM electrode characterization

We characterized our icFM microelectrode response (Vrms) for different electrolyte concentrations (1X - 10X PBS) at various AC source frequencies (0.1 – 10 MHz) and peak-to-peak source voltages (0.2 - 2 Vp-p) to characterize the MIC electrode performance with the applied voltage, source frequency and electrolyte concentration. Ideally, the differential voltage (Vrms) between the measuring electrodes should be zero when placed identically on either sides of the source electrode under similar electrochemical conditions. However, the non-zero differential voltage observed in real-world MIC setups arises from the asymmetry in the electronic system and the system noises and offsets. We relied on the continuity and linearity of this differential signal with changes in operating conditions to evaluate our electrode stability since breakdown in electrode operation would result in a discontinuous response. First, we measured the mean Vrms response of the electrodes for electrolyte concentrations from 1X to 10X PBS over a broad frequency range (0.5-10 MHz) at 2 Vp-p. We observed a continuous and monotonically increasing Vrms as a function of electrolyte concentration which suggests that the electrodes are not undergoing electrolysis or corrosion at the high saline concentrations used here (Zeitler et al. 1997) (**Figure 3a**). Importantly, we observed a linear Vrms output with increasing Vp-p (0- 2 V for 10X PBS) which confirms that these electrodes are electrochemically stable up to 2 Vp-p (**Figure 3b**). It is likely that the linearity can be extrapolated to even higher source voltages suggesting a wider operable range.

**Figure 3:**
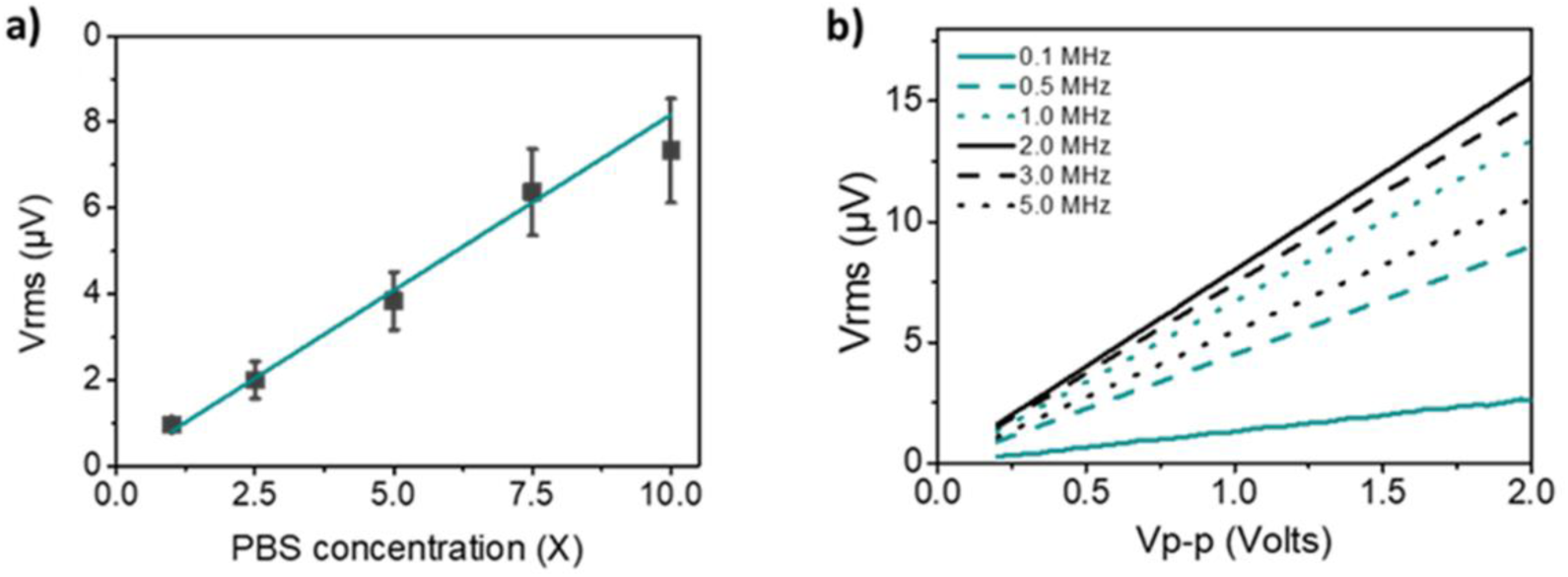
Electrode stability: a) Output voltage (Vrms) measurement for coplanar icFM electrode as a function of electrolyte (PBS) concentration averaged over different source frequencies (0.1 - 10 MHz) at 2 Vp-p is plotted. b) Plot of Vrms for a voltage sweep from 0.2 to 2 Vp-p source voltage performed at various frequencies (0.1 - 5MHz) at 10X PBS is displayed.

### 3.2 RBC detection with icFM electrode

To demonstrate the icFM microelectrode’s compatibility with MIC, we detected human erythrocytes (or RBC). A suction pressure of 10 kPa maintained the sample at a mean velocity of **∼** 15 mm/sec over the electrodes that was optimized for a sampling rate of 7200 Hz. The differential impedance signature of an erythrocyte in the electrolyte as it flows over the microelectrode array is characteristic of a well-described single ‘sine-wave’ like shape in this configuration (**Figure 4a**) (Gawad et al. 2001). The observed variability in signal peak distribution arises due to the variable distance of the flowing cells from the electrodes and inherent cell size dispersity.

**Figure 4:**
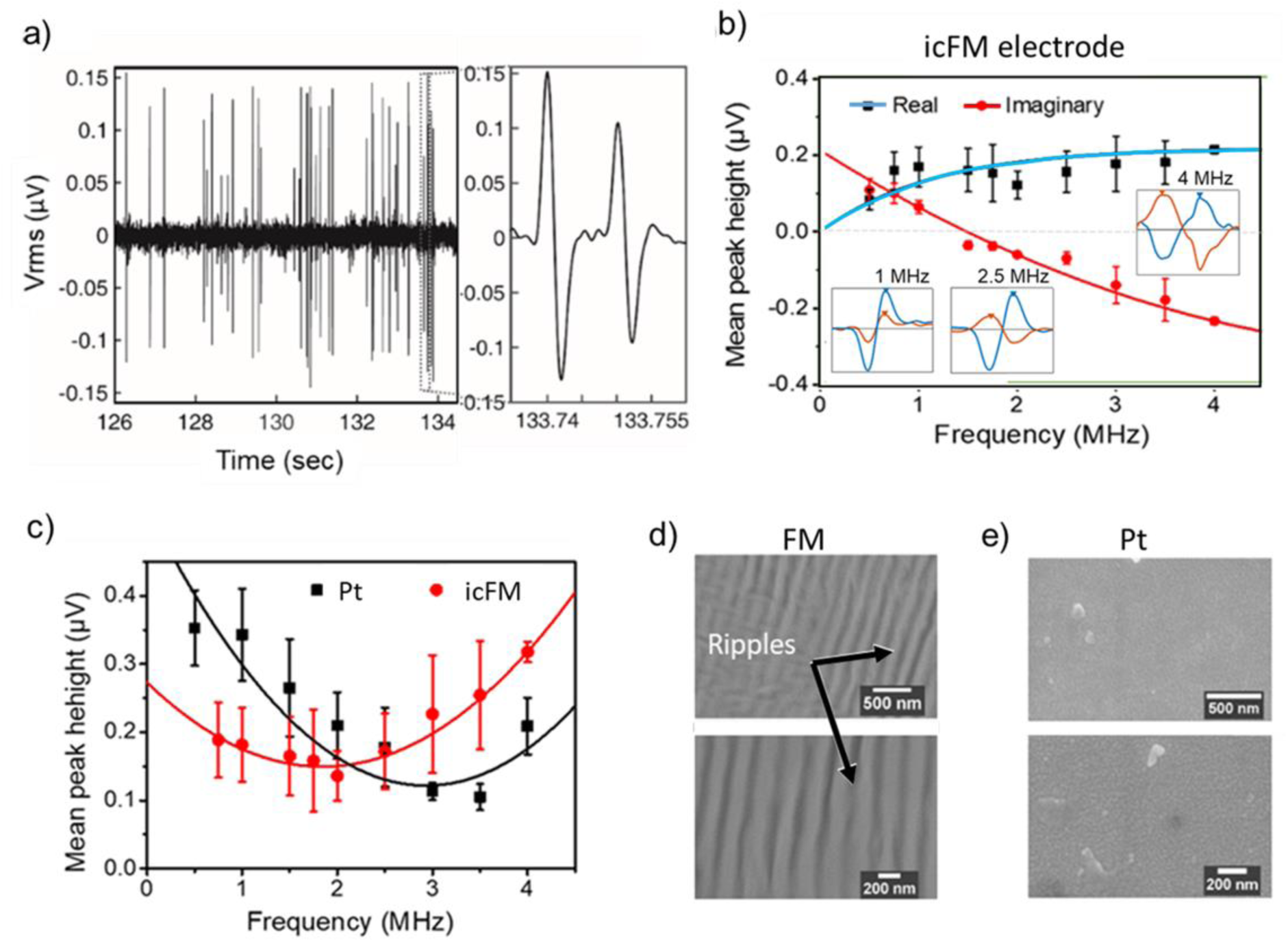
Single erythrocyte detection with coplanar icFM microelectrode a) Differential voltage signal from the icFM MIC setup at 2 Vp-p and 0.5 MHz for flowing dilute suspension of RBC in 10X PBS is shown. (Inset) Each positive-negative peak pair is representative a single RBC flowing over the electrodes. b) Real and imaginary component of the voltage signal after lock-in amplification over a frequency range of 100 kHz to 4 MHz is shown. (Inset) Real and imaginary component of individual signals for RBC detection at represented frequencies is shown. c) Mean peak heights of differential voltage signal for RBC detection obtained using icFM and platinum electrodes at different frequencies at 2Vp-p source voltage is shown. d) SEM micrographs of FM and Pt electrode surfaces at two magnifications is shown.

We further analyzed the real (in-phase) and imaginary (out-of-phase) components of the voltage signal obtained after the lock-in amplification (**Figure 4b**). The highest frequency tested was limited to 4 MHz due to the low signal to noise ratio at higher frequencies as is evident from the irregular shape of erythrocyte signal at 4 MHz. We observed that the imaginary component of the average signal (from ∼ 400 cells per measurement) changes from positive to negative values at ∼1.5 MHz as a result of the varying complex permittivity of the system. The permittivity of the cell and suspending medium at this frequency is governed by the ‘first relaxation’ that occurs due to the polarization of the cell membrane-medium interface. The observation is in agreement with the model for impedance spectra of cells as described by Maxwell’s mixture theory for cells in suspension (Morgan et al. 2006; Sun and Morgan 2010). This frequency response is also a function of the biophysical and morphological properties of the cell (and subcellular components like vacuoles and nucleus) and has been used previously for cell classification on the basis of size, cytoplasmic resistance and membrane capacitance (Sun and Morgan 2010).

### 3.3 Comparison between icFM and Pt electrode

A comparison of icFM and the widely used Pt electrodes in equivalent microchannel dimensions demonstrates that the average peak signal values registered by icFM electrodes are comparable to Pt electrodes over the frequency range of 100 kHz to 4 Mhz (**Figure 4c**). We observed a higher signal at low frequencies (< 2 MHz) from Pt electrode along with marginally higher noise from icFM at higher frequencies (> 3Mhz). This frequency dependent variation can be attributed to the effect of surface roughness on impedance of the system. SEM micrographs of electrode surfaces shows comparatively rough and rippled surface of solidified FM as compared to the smooth surface of sputtered Pt electrode (**Figure 4d**). We speculate that the ripples on FM surface may be arising due to the compressive stresses along the surface during solidification. The higher roughness provides a higher effective surface and thus a higher double layer capacitance that reduces the overall impedance of the system at low frequencies and attenuates the signal strength (Morgan et al. 2006). Nevertheless, the signals from both the electrodes are strong enough for cell detection within the accounted frequency range. Therefore, we establish that the icFM electrodes can serve as a viable alternative for MIC based single cell detection.

### 3.4 Water-in-oil droplet detection

To test the electrodes for multiple phase fluid operations, we employed icFM electrodes to detect water-in-oil droplets (Teh et al. 2008). Use of icFM microelectrodes allowed us to have PDMS walls on all four sides and thus, the microchannels in our devices exhibit uniform hydrophobicity which aids droplet microfluidic operations (Subramanian et al. 2011). Droplets of 1X PBS solution were generated using a T-junction upstream of the microelectrodes. The signal for aqueous droplets is characteristic of a high dielectric feature in a continuous oil phase flowing through the channel (**Figure 5a**). Comparison of the distribution of peak heights and corresponding full width at half maxima (FWHM) for the erythrocytes and droplets in oil is consistent with the two flow profiles (**Figure 5b**). A large standard deviation of the FWHM and peak heights for the RBC (12 % and 25% respectively) results from the variable velocities due to parabolic flow profile in the microchannel and the variable normal distance of the cells from the electrodes respectively. On the other hand, droplets show significantly narrow distribution of peak heights (2 %) and FWHM (2 %) as a result of the plug flow profile of the droplets. The distribution in the droplet signal also agrees well with the polydispersity of the droplet plug length (1.7 %) as calculated via imaging (**Figure 5c**). Therefore, our MIC devices can robustly detect particle and droplet features. It is known that electrolysis and corrosion may degrade the electrodes over time with continuous usage in saline conditions (Zeitler et al. 1997). Interestingly, we observed an excellent overlap in the primary peak height distribution for the droplets even after continuous measurements on the same device over time (up to 60 mins) (**Figure 5d**). This suggests long-term stability of the icFM electrodes in such two phase microfluidic system.

**Figure 5:**
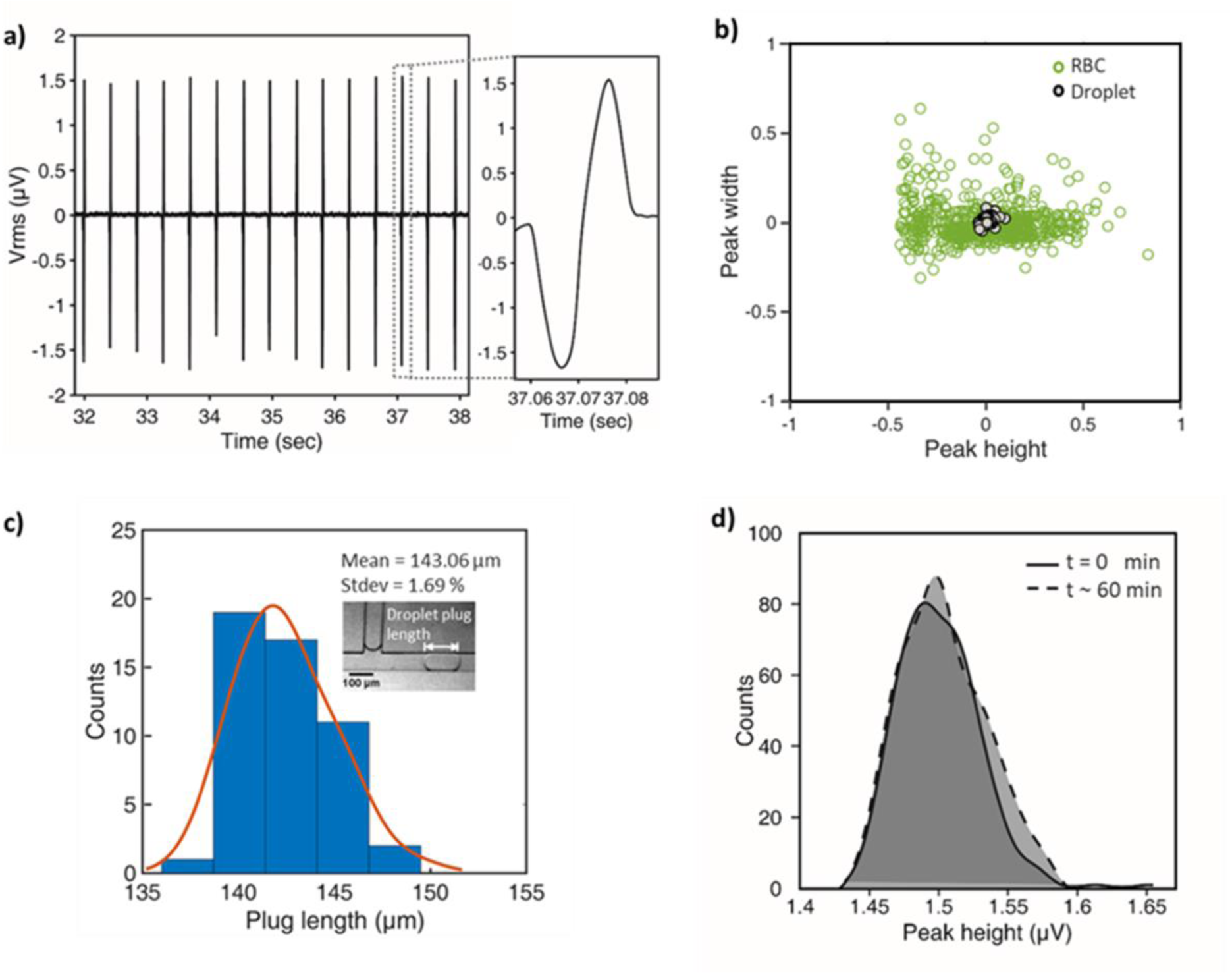
Water-in-oil droplet detection with icFM microelectrode **a)** Differential voltage signal from MIC setup for droplet detection at 2 Vp-p and 1 MHz for droplets flowing across the electrodes is plotted. (Inset) A zoomed view of the signature waveform for a single droplet detection is displayed. **b)** Comparison of peak height and corresponding FWHM distribution obtained at 2 Vp-p and 1 MHz for RBC and droplet signals is plotted **c)** Optically measured droplet plug-length distribution with 18 kPa suction pressure across the flow channel is displayed (red curve is a guassian fit). (Inset) Optical image of droplet and its position where the plug length is measured. d) Peak height distribution for droplet signals measured **∼** 60 minutes apart on the same device is shown.

### 3.5 Cell-in-droplet quantification

Cell entrapment in droplets is an increasingly popular single cell analysis tool which currently relies upon optical imaging for quantification (Guo et al. 2012; Joensson and Andersson Svahn 2012; Lu et al. 2017). Since imaging speeds can be limiting and sometimes require labeling, we used our icFM integrated MIC setup to quantify cells entrapped in droplets. In a device described above, we generated droplets of 1X PBS with suspended RBCs at a dilution of ∼ 10^7^ RBCs per ml to ensure close to single occupancy of cells in droplets (**Figure 6a**). We hypothesized that an additional set of secondary signatures superimposed on the droplet signal will be discernible if the droplet contained cell(s) owing to their membrane capacitance (**Figure 6b**). Since the cytoplasm is shielded by the low dielectric of the cell membrane at frequencies below 10 MHz, its cytoplasmic conductivity does not contribute to our impedance measurements. Therefore, the cell-in-droplet signal would be defined by a low dielectric feature (cell) suspended in a high dielectric medium (droplet) as its travels in a low dielectric medium (oil). Thus, cells-in-droplets would display a superimposition of the cell signature on the droplet signal resulting in primary peak (from droplet) and a trough (from cell) that eventually forms a secondary peak (**Figure 6b** and **Supplementary figure S2**).

**Figure 6:**
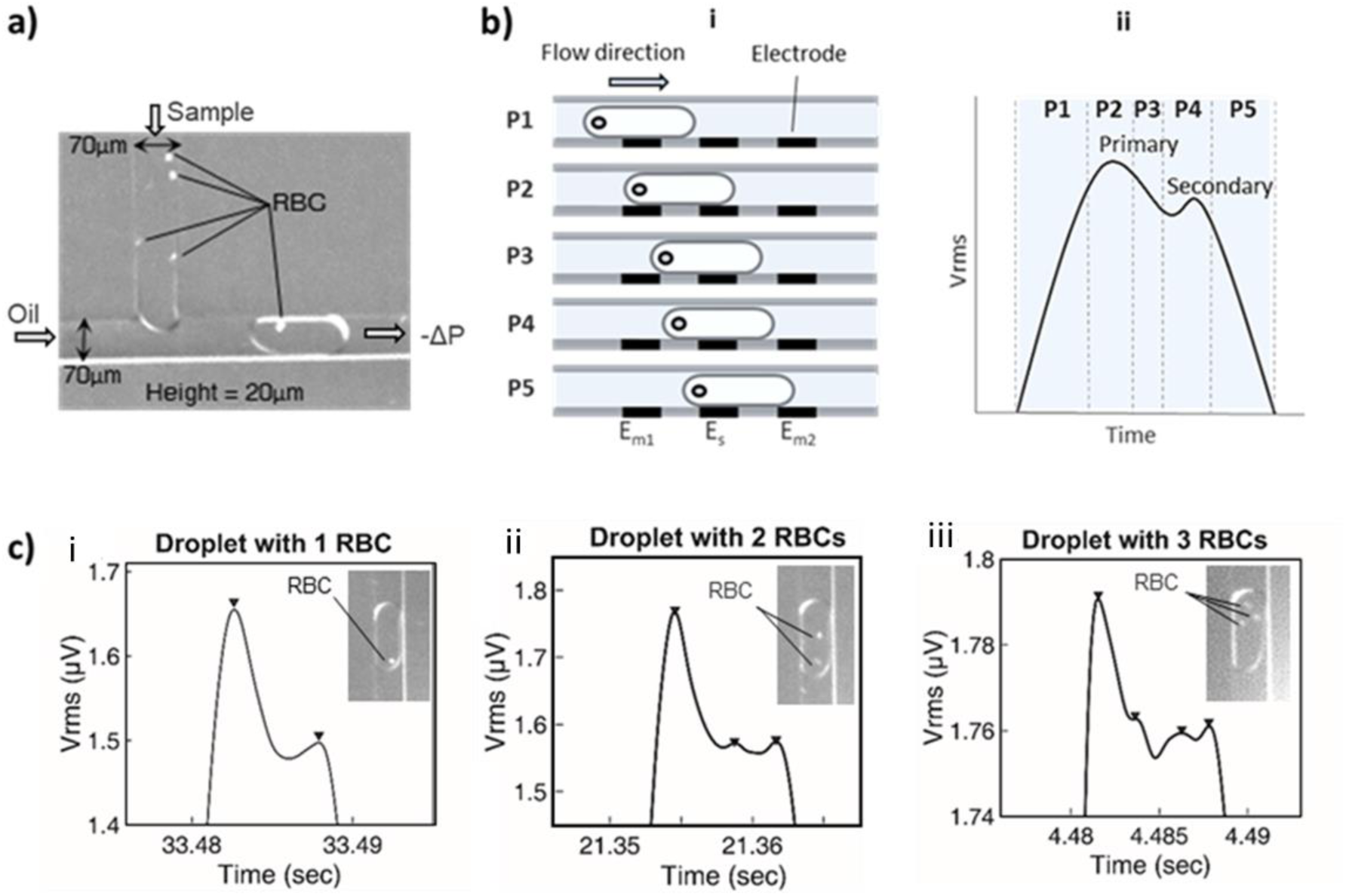
Cell-in-microdroplet quantification **a)** T-junction microfluidic channel used to encapsulate cell(s) in droplets upstream of the microelectrode array is shown. **b)** Schematic representing evolution of primary and secondary peaks in MIC setup for droplet and cell-in-droplet, respectively is depicted **i:** Droplet (containing a single cell) at specific positions relative to the microelectrode array (E_m1_, E_m2_: measuring electrodes, E_s_: source electrode) is pictorially shown. **ii:** Contribution of each position to differential signal is schematically plotted (only positive peak of differential signal is displayed here, the negative peak should display similar features). **c)** Differential voltage signal at 2 Vp-p and 1 MHz for droplets carrying cells **i-iii:** Primary peak representing a droplet and secondary peaks representing number of entrapped RBCs is plotted. (Inset) Optical image of different droplets containing as many RBCs is shown.

The primary and the secondary peak reach their corresponding maxima when the contributor (droplet and cell in droplet, respectively) passes over the center of the source and measuring electrode. Multiple cells in a droplet can be similarly seen as multiple secondary peaks. Here, we assumed superimposition of peaks from entrapped cells due to physical overlap is a low likelihood event and should not contribute to large discrepancies in our peak based analysis.

The MIC signal for RBCs in droplets indeed displayed secondary peaks consistent with the average number of cells loaded per droplet (**Figure 6c**). We could identify up to three discrete secondary peaks representing droplet entrapping zero to three RBCs for our cell suspension. To validate our results, we compared the fraction of droplets with one or more RBC counts from the number of secondary peaks at different dilutions of our erythrocyte suspension with the RBC counts obtained optically (**Figure 7**). With increasing concentration of the cells, the fraction of the droplets with increasing numbers of entrapped cells goes up as expected (**Figure 7a**). We find a good agreement between the two methods as seen in **Figure 7b**. This demonstrates that our icFM microelectrodes are compatible and sensitive enough to detect single cells and particles in complex two-phase microfluidic systems.

**Figure 7:**
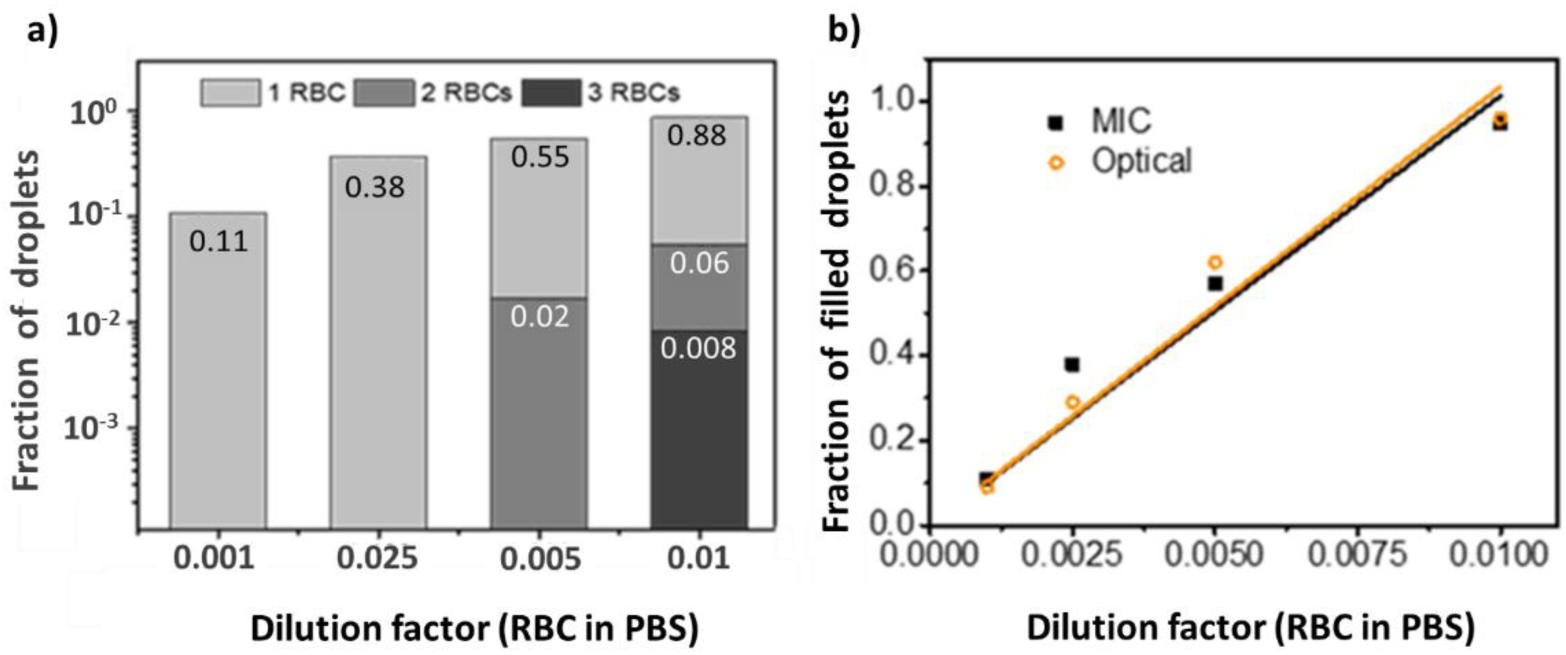
RBC count analysis **a)** Fraction of droplets encapsulating one, two and three RBCs at represented dilutions is plotted. **b)** Comparison between RBC count at representative dilutions as measured optically (orange) and MIC (black) is plotted.

## 4. Conclusions

We have demonstrated Field’s metal based ‘in-contact’ (icFM) microelectrodes as a rapid and economical alternative for microfluidic impedance cytometry that is capable of single cell quantification. We present a technically simple fabrication process that produces highly reproducible microelectrodes in a cleanroom-free environment. Our microelectrodes are robust in all solution conditions used for conventional cell analysis and comparable in performance to state-of-art Pt electrodes. This allows us to monitor cells in water-in-oil droplets non-optically with high sensitivity. In the future, incorporation of non-conductive flow focusing or sheath flow to reduce the detection volume can be further employed with icFM devices to achieve a higher sensitivity (Bernabini et al. 2011; Rodriguez-Trujillo et al. 2007). Two opposing icFM electrode layers sandwiching a flow layer in 3D electrode geometry can further improve the MIC signal strength, thus enabling more challenging applications like cell classification and microorganism detection.

## Acknowledgements

This project was partially funded by support from the Indian Institute of Science (IISc) Bangalore, DBT Biodesign Bioengineering Initiative and Rao Biomedical Research Fund. We acknowledge MicroX Labs and Logic-fruit technologies for technical discussions and support. We also acknowledge use of the lithography facilities at the Center for Nano Science and Engineering (CeNSe), funded by the Department of Information Technology, Gov. of India. We thank Prosenjit Sen and Karthik Mahesh, (CeNSe) for providing help and access to their MIC setup; Abhishek Ranade for SEM imaging; Priyanka V. and Satyaghosh Maurya for their support in device fabrication and data analysis, respectively and Lakshmi Supriya, Prithiv Natarajan and Suraj Jagtap for their inputs on the manuscript.

