## Supplementary figure S1 for "Integrated Field’s metal microelectrodes based microfluidic impedance cytometry for cell-in-droplet quantification"

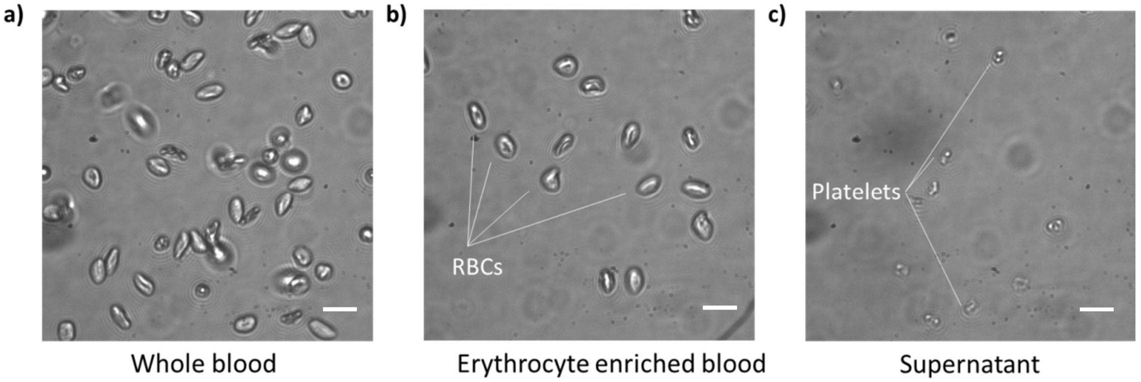


**Figure S1:** Erythrocyte enrichment. Brightfield microscopy images of **a)** Human whole blood. **b)** Sample after erythrocyte enrichment. **c)** Discarded supernatant. Scale bar is 5 microns.
