## Supplementary table T1 for "Integrated Field’s metal microelectrodes based microfluidic impedance cytometry for cell-in-droplet quantification"

| | Pt Electrodes | FM Electrodes |
|---|---|---|
| | Time (mins) | Time (mins) |
| **Process (clean-room)** | | |
| FNA cleaning and $N_2$ drying of wafer | 10 | Nil |
| Dehydration at 230 °C | 10 | Nil |
| Pt Sputtering | 30 | Nil |
| IPA cleaning and $N_2$ drying | 10 | 10 |
| Dehydration at 230 °C | 120 | 10 |
| Photolithography | 30 | 30 |
| IBM etching | 30 | Nil |
| Final cleaning (NPM sonication/$O_2$ Descum) | 30 | Nil |
| **Process (non clean-room)** | | |
| Soft lithography | 90 | 90 |
| Metal filling | Nil | 10 |
| Solidification | Nil | 10 |
| Plasma bonding | 10 | 10 |
| Clean-room time | 270 | 50 |
| Non clean-room time | 100 | 120 |
| Total time | 370 | 170 |
| **Requirement for 6n devices with electrodes (assuming 6 electrode pairs each wafer)** | | |
| Clean-room requirement | n X 270 | 60 (one time) |
| Total time | n X 370 | 60 + n X 120 |

**Table T1:** A comparison between FM and conventional Pt microelectrode fabrication processes in regards to fabrication time.
