## Supplementary figure S2 for "Integrated Field’s metal microelectrodes based microfluidic impedance cytometry for cell-in-droplet quantification"

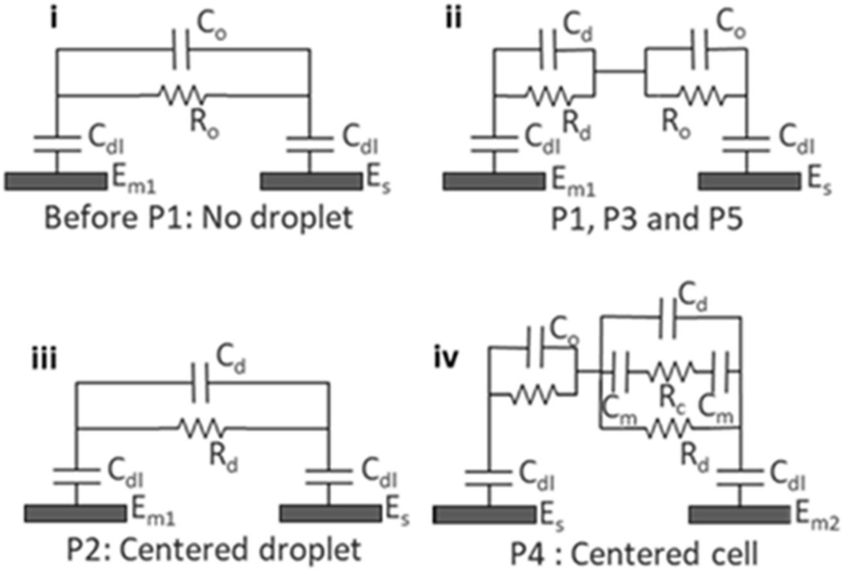


**Figure S2:** Equivalent electrical circuit model for the droplets at different positions is drawn ($C_{dl}$: equivalent double layer capacitance due to all interfaces (electrode-oil, oil-electrolyte and electrolyte-electrode); $C_o$, $R_o$, $C_d$, $R_d$: capacitance and resistance due to oil and droplet respectively; $C_m$: cell membrane capacitance, $R_c$: cytoplasmic resistance) **i:** Base impedance without droplet, accounting for low conductivity and dielectric constant of carrier phase (Oil). **ii:** As the aqueous droplet touches $E_{m1}$ (and not $E_s$), the higher dielectric constant of droplet starts to contribute to the signal. **iii:** Primary peak appears as the droplet touches both the electrodes reaching maximum conductivity and dielectric constant between the electrodes. **iv:** Secondary peak appears as the cell crosses the centre of $E_s$ and $E_{m1}$.
